# Thal-Kak: unifying biomolecular structure predictors reveals a sampling–selection gap

**DOI:** 10.64898/2026.08.19.745680

**Authors:** Junhyeok Bae, Soohyun Jo, Yeajin Kim, Dongyun Kim, Kwanwook Kim, Sanghyun Park, Sanggeun Park, Sojung Myung, Hyunho Shin, Min Hyeok Kim, Minji Kang, Minkyung Baek

## Abstract

Complementary all-atom structure predictors sample different solutions, but how to allocate a fixed sampling budget across them and select the best output remains unclear. Thal-Kak unifies five released predictors under shared upstream inputs and a common schema. Across FoldBench and CASP16, model mixing improves oracle sampling over single-model runs, but selection remains a bottleneck because confidence scores do not transfer across models and existing quality-assessment methods cannot resolve this gap.

## Main

Biomolecular structure prediction is now served by several all-atom predictors, including AlphaFold3 (AF3)^1^, Boltz-2^2^, Chai-1^3^, Protenix v1/v2^4,5^, and ESMFold2^6^. These models are complementary^7^, but exploiting their collective strength remains difficult because they rely on different software environments, alignment pipelines, and input formats. As a result, it remains unclear whether a fixed sampling budget is better spent on one predictor or distributed across several.

We built Thal-Kak to run multiple predictors from a shared assembly description and multiple sequence alignment (MSA) under a common input and output schema (Fig. 1a). Thal-Kak supports Boltz-2, Chai-1, Protenix v1/v2, and ESMFold2, and is available as open-source software at https://github.com/CSSB-SNU/Thal-Kak and through a Google Colab notebook. AF3 is used only as an external benchmark because its model parameters are subject to separate licence terms. By relying on openly released predictors, Thal-Kak also enables multi-model sampling in settings that require commercial use or fewer downstream-use restrictions. All comparisons used ColabFold-API alignment^8^ unless noted otherwise. These did not measurably reduce accuracy relative to native alignment pipelines on the benchmarks tested (Supplementary Fig. 1 and Supplementary Table 1).

**Figure 1.**
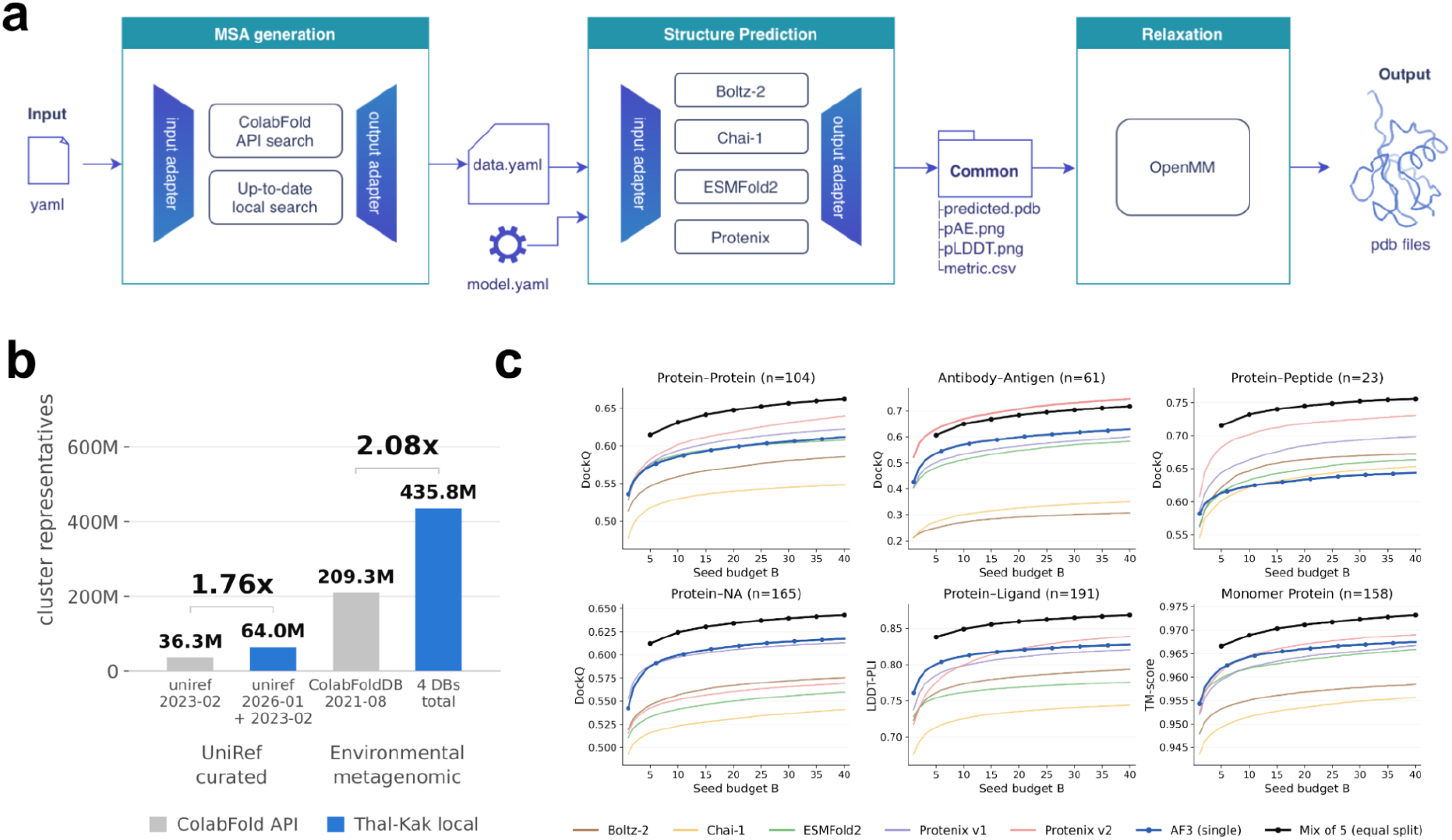
Thal-Kak and the sampling gain from combining released predictors. **a**, Overview of the Thal-Kak workflow. A shared MSA and template input is generated using either the ColabFold API or the Thal-Kak local search, converted to a common format, and passed to Boltz-2, Chai-1, Protenix v1/v2, and ESMFold2 through predictor-specific adapters. Outputs are collected in a common format and selected structures can undergo restrained relaxation. **b**, Number of cluster-representative sequences available to the Thal-Kak local mmseqs search and ColabFold API. Thal-Kak provides 64.0 million UniRef representatives versus 36.3 million in ColabFold, and 435.8 million environmental and metagenomic representatives versus 209.3 million in ColabFoldDB. **c**, Oracle sampling performance across six FoldBench target classes as a function of total seed budget (B). The AF3 baseline is shown in blue, the equal-split mixture of the five released predictors in black, and individual released predictors as lighter curves. For the five-model mixture, (B/5) seeds are assigned to each predictor. Performance is measured by DockQ for protein-protein, antibody-antigen, protein-peptide, and protein-NA targets, lDDT-PLI for protein-ligand targets, and TM-score for protein monomers. Curves show mean performance across targets and repeated random seed subsampling. The five-model mixture generally outperforms AF3 at matched seed budgets, except for antibody-antigen targets, where Protenix v2 performs best alone.

Thal-Kak also provides a local search against larger and more recent sequence databases than those available through the ColabFold API (Fig. 1b). The local search combines complementary MMseqs2^9^ and HHblits^10^ searches and produces deeper alignments with higher Neff. These additional sequences improve prediction accuracy for a subset of the monomer and protein-multimer targets in FoldBench^7^ (Supplementary Fig. 2). Thal-Kak also maintains an up-to-date local template database for practical use, while benchmark searches apply a temporal cutoff to avoid using templates unavailable before the corresponding target.

We benchmarked the five released models across six FoldBench target classes, with each seed generating five candidate structures. For mixed-model pools, the total seed budget was divided equally among the participating predictors. The five-model mixture generally improved oracle sampling over any single predictor, including AF3, and approached the performance of a 40-seed AF3 run with substantially fewer seeds (Fig. 1c). Because each predictor contributed only a fraction of the total seeds, the gain reflects complementary sampling across models rather than deeper sampling of any one predictor. Antibody-antigen targets were the exception, for which Protenix v2 performed best alone. The same overall trend was observed on CASP16 targets (Supplementary Fig. 3a).

Oracle performance, however, measures the best structure present in the sampled pool. In practice, a user must select a single candidate without access to its ground-truth accuracy. Even within individual predictors, the structure ranked highest by native confidence was often substantially worse than the oracle, with the largest gap observed for antibody-antigen targets (Fig. 2a). Model mixing increased oracle performance but did not improve Top-1 performance because predictor-native confidence scores are not directly comparable across models (Supplementary Fig. 4). A high-quality structure from one predictor can therefore be outranked by a less accurate structure from another. Consequently, mixed pools did not outperform the best individual predictor under native-confidence ranking despite sampling better structures more frequently (Fig. 2a,b and Supplementary Fig. 3b). Improved sampling therefore exposes selection as a major remaining bottleneck.

**Figure 2.**
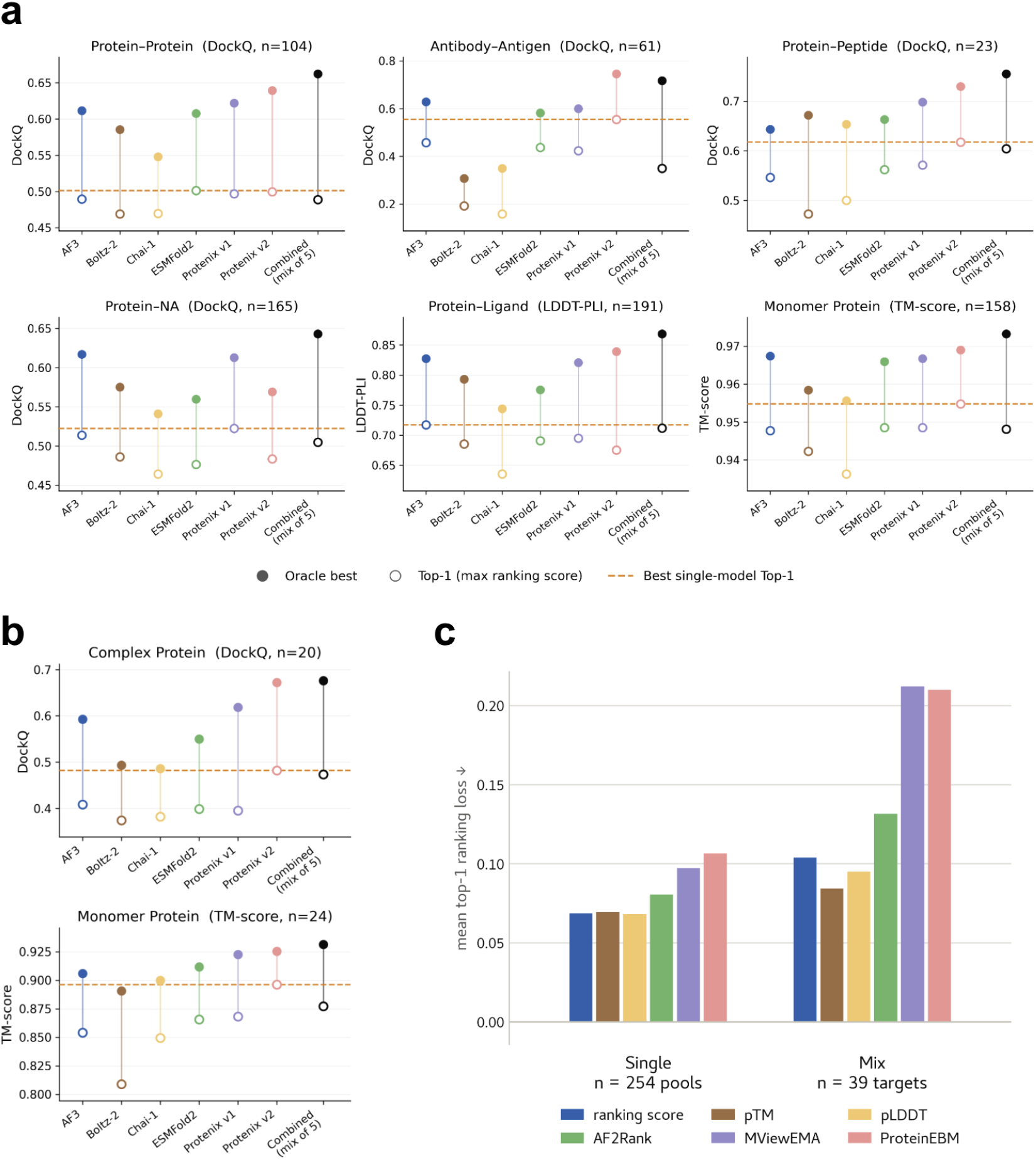
Selection limits the benefit of improved sampling. **a**, Oracle and Top-1 performance across six FoldBench target classes at a fixed budget of 40 seeds. Filled circles indicate the best sampled structure in each pool, and open circles indicate the structure selected by the predictor-native ranking score. Single predictors use 40 seeds, whereas the five-model mixture uses 8 seeds per predictor. The dashed line marks the best Top-1 performance among individual predictors. Metrics are DockQ for protein-protein, antibody-antigen, protein-peptide, and protein-NA targets, lDDT-PLI for protein-ligand targets, and TM-score for protein monomers. **b**, Corresponding analysis on CASP16 protein complex and monomer targets. **c**, Quality-assessment performance on CASP16 using predictor-native ranking score, pTM, and pLDDT, and the dedicated scorers AF2Rank, MViewEMA, and ProteinEBM. Bars show mean Top-1 ranking loss, with lower values indicating better selection, for candidates from individual predictors (Single, n=254 pools) or pooled across predictors (Mix, n=39 targets). Dedicated quality-assessment methods do not improve selection over predictor-native confidence and degrade further when predictions from different models are pooled.

Dedicated quality-assessment methods did not resolve this problem. AF2Rank^11^, ProteinEBM^12^, and MViewEMA^13^ failed to consistently identify the best protein structures and performed particularly poorly when predictions from multiple models were pooled (Fig. 2c). Native confidence generally remained the stronger selector, indicating that robust ranking across heterogeneous prediction pools remains an open problem.

A selector that generalizes across predictors may require training data spanning both a broad range of structural accuracy and diverse modes of prediction error. Multi-predictor sampling naturally generates such heterogeneous candidate pools, including near-native structures and substantially incorrect conformations produced by models with different biases. These pools may therefore provide useful training data for model-agnostic quality assessment. Thal-Kak serves not only as a unified prediction workflow, but also as a platform for generating such structural decoys at scale.

Together, these results show that predictor diversity can substantially improve sampling while shifting the remaining challenge toward selection. We release Thal-Kak together with its heterogeneous prediction pools and per-structure accuracy annotations to support the development and evaluation of selection methods that generalize across predictors.

## Methods

### The Thal-Kak pipeline

Thal-Kak provides a unified interface for running multiple biomolecular structure predictors. A standard run comprises three stages: MSA and template search, structure prediction, and restrained relaxation. The alignment and template search is performed once for each target and reused across the selected predictors. Each predictor then converts these shared upstream inputs into its native representation and generates candidate structures using its own inference procedure. Candidate structures are ranked using the predictor’s native confidence score, and the five highest-ranking valid structures are passed to a common relaxation protocol. This design controls the upstream alignment and downstream processing across predictors while retaining each predictor’s native featurization and inference procedure.

The released pipeline supports Boltz-2, Chai-1, Protenix v1, Protenix v2, and ESMFold2. AlphaFold3 is evaluated separately as an external benchmark but is not distributed as part of Thal-Kak because its model parameters are governed by separate licence terms. Thal-Kak is available as open-source software at https://github.com/CSSB-SNU/Thal-Kak and as a Google Colab notebook for users without local computing resources. The input format and notebook interface are described in Supplementary Methods S1.

### Alignments and databases

Thal-Kak obtains protein MSAs and templates using either the ColabFold MMseqs2 API or a local search pipeline. Both modes produce paired and unpaired per-chain alignments and template mmCIF files in a common representation for downstream structure prediction.

In ColabFold mode, each distinct protein sequence is submitted to the ColabFold MMseqs2 API server. For multichain targets, the returned complex alignment is separated into paired and unpaired alignments for each sequence. Template hits are ranked by E-value, and the four highest-ranking templates per chain are retained. If remote template retrieval fails, the search is repeated in alignment-only mode and prediction proceeds without templates.

RNA MSAs are generated locally using nhmmer^14^ searches against clustered Rfam^15^ and RNAcentral^16^ databases. DNA chains are provided to the prediction models by sequence alone.

The local search for proteins uses UniRef30^17^, UniRef100^18^, MGnify^19^, Logan^20^, BFD^21^, and EnVhog^22^. Protein sequences can be searched using MMseqs2, HHblits, or a combination of the two. The MMseqs2 and HHblits workflow uses UniRef-derived profiles to search the environmental sequence collections. When both engines are used, their alignments are merged, deduplicated, and filtered by sequence identity and coverage. For multichain targets, taxonomic pairing is performed using UniRef30, which provides the taxonomy annotations required for interchain pairing. The paired alignment is capped at 8,192 records per chain, and total alignment depth is capped at 16,384 records including the query.

Local template searches use a February 2026 snapshot of the Protein Data Bank^23^. An optional release-date cutoff can be applied to exclude templates unavailable before a specified date. Details of the local MSA/template database and search procedure are provided in Supplementary Methods S2.

### Structure prediction and sampling

Each predictor receives the same target description, including the available alignments, templates, ligands, and assembly stoichiometry, subject to the input features supported by that backend. Predictor-specific runners convert this shared representation into the native input format required by each model.

Predictions are generated across a contiguous range of random seeds using each backend’s native stochastic sampling procedure. Each seed produces five candidate structures. For each candidate, Thal-Kak collects the confidence measures reported by the corresponding predictor into a common-schema table. These include mean pLDDT, predicted TM-score, interface predicted TM-score, and the predictor’s composite ranking score where available. Backend-native outputs are retained alongside the standardized outputs.

Unless otherwise stated, the predictor-native ranking score is used to rank candidate structures because neither external quality-assessment models nor any single confidence score consistently outperformed the alternatives on the FoldBench (Supplementary Fig. 5) and CASP16 QA benchmark (Fig. 2c). Before relaxation, candidates are screened for polymer bond connectivity, and the five highest-ranking valid structures are retained. Predictor-specific input handling, MSA subsampling, polymer bond connectivity criteria, and inference configurations are described in Supplementary Methods S1 and S3.

### Relaxation

Selected structures are subjected to restrained all-atom energy minimization using OpenMM 8.4.0^24^ and PDBFixer 1.12.0^25^ with a modified AlphaFold2^21^ relaxation procedure. The system is represented using OBC2 implicit solvent^26^. The force-field stack includes amber19-all with ff19SB for proteins^27^, OL21 for DNA^28^, OL3 for RNA^29^, GLYCAM_06j-1 for carbohydrates^30^, monatomic-ion parameters, and GAFF 2.11 with AM1-BCC charges for small-molecule ligands^31,32^. Non-hydrogen atoms are restrained relative to their starting coordinates using confidence-dependent flat-bottom harmonic potentials. Pre-relaxation stereochemical validation, force-field details, and the complete minimization procedure are described in Supplementary Methods S4.

### Benchmarks

We benchmarked the five predictors distributed with Thal-Kak and included AlphaFold3 as an external baseline. To harmonize upstream inputs across predictors and maintain consistency with the FoldBench evaluation framework, all benchmark predictions used protein alignments generated through the ColabFold API workflow. Templates were restricted to structures available on or before 31 December 2023. RNA alignments were generated using the Thal-Kak local RNA search pipeline.

#### Target selection and temporal filtering

FoldBench bioassemblies released after 1 January 2024 were retained (681 structures) to ensure post-training evaluation relative to the predictors’ 2023 training cutoff. CASP16 targets were taken as provided in the official experiment (101 entries with experimentally resolved structures), as the dataset already satisfies temporal separation from the training data.

#### Input preparation

FoldBench targets were prepared following the published FoldBench procedure. Reference mmCIF files were converted using the AlphaFold3 Input.from_mmcif routine from commit 948827f. bondedAtomPairs records and crystallization aids listed in AlphaFold3 Supplementary Table 9 were removed^1^. Target sequences were subsequently written to FASTA files. Covalently linked glycans that would otherwise require multiple Chemical Component Dictionary entries within a single chain were represented as SMILES strings to provide a common input representation across predictors.

#### Inference configuration

The predictors used in this study were AlphaFold3 at commit 1e9ece2, Boltz-2 at commit cb04aec, Chai-1 at commit af596cb, Protenix v1/v2 at commit c3bfc36, and ESMFold2 at commit fba1e42. For each predictor and target, 40 random seeds were generated with five candidate structures per seed, giving up to 200 candidates per predictor. Inference used 10 trunk-module recycles and 200 diffusion steps where these parameters were supported by the corresponding backend. ESMFold2 could not process a subset of FoldBench targets substantially longer than 1,000 residues because of GPU memory limitations.

#### Structural evaluation

Chain correspondences between reference and predicted structures were established using the US-align implementation in the compare-structure command of OpenStructure v2.11.1^33^. Reference structures were decomposed into the chain pairs defined by FoldBench, and predicted structures were decomposed according to the resulting mapping. For protein-ligand targets, protein chains were mapped using OpenStructure while the remaining ligand residues were retained. For protein monomer targets, predicted residues were mapped to the reference sequence using Needleman-Wunsch alignment^34^ in Biopython^35^.

DockQ and ligand RMSD were calculated with DockQ v2.1.3^36^ using the chain mapping above. Modified polymer residues were handled with a custom correction to prevent their misclassification as small molecules by DockQ, as described in Supplementary Methods S5. For protein-ligand targets, lDDT-PLI^37^ and lDDT-LP^37^ were calculated using the compare-ligand-structures command in OpenStructure v2.11.1. For protein monomers, TM-score was calculated using TMscore^38^.

#### Seed-budget allocation

Each point on the seed-budget axis corresponds to a fixed total number of prediction seeds, denoted by (B). For a single predictor, (B) seeds were sampled at random from its 40 available seeds, producing (5B) candidate structures. For a mixed pool containing (M) predictors, the same total budget (B) was divided equally among the participating predictors, with each contributing (B/M) randomly selected seeds. Mixed-model analyses were therefore restricted to values of (B) that were integer multiples of (M).

For each candidate pool, the oracle candidate was defined as the structure with the best ground-truth accuracy metric. The Top-1 candidate was defined as the structure with the highest predictor-native ranking score. Ranking scores were used as reported and were not normalized across predictors. Ties were resolved by random selection. The ground-truth accuracy of the selected candidate was averaged across targets. Random seed subsampling was repeated 100 times for each seed budget, and the mean across repetitions was reported.

#### Quality-assessment scorers

We evaluated AF2Rank, MViewEMA, and ProteinEBM as dedicated protein structure quality-assessment methods. AF2Rank was reproduced using ColabDesign 1.1.1 with AlphaFold2 model_1_ptm parameters in fixed-backbone template mode with one recycle. Template sequences and side chains were masked while interchain contacts were preserved. The AF2Rank score was calculated from pTM, pLDDT, and the TM-score between the input and output structures. Multichain targets were processed by the monomer model as concatenated chains. MViewEMA was run using the authors’ released Singularity image and checkpoint with default settings. ProteinEBM was run using the second-release model_6_expert_frozen_1m_md checkpoint in the expert-model configuration with template self-conditioning. ProteinEBM energy values were sign-inverted so that higher values indicated better predicted quality.

#### Quality-assessment comparison on CASP16

We compared three predictor-native confidence measures, ranking score, pTM, and pLDDT, with AF2Rank, MViewEMA, and ProteinEBM. Targets containing nucleic acids were excluded because the dedicated quality-assessment methods were evaluated only on protein structures. The initial candidate pool contained up to 200 structures per target from each of six predictors: AlphaFold3, Boltz-2, Chai-1, ESMFold2, Protenix v1, and Protenix v2. For ESMFold2, which emits no ranking score, the ranking score was defined as 0.8·ipTM + 0.2·pTM following the AlphaFold3 convention, and pLDDT was rescaled from 0–1 to 0–100. Ground-truth accuracy was evaluated using TM-score for monomers and DockQ for multimeric interfaces. To compare scoring methods on identical candidate sets, predictor-target pools were excluded if any scoring method failed. Selection was evaluated in two settings. In the single setting, each method ranked candidates from one predictor at a time, resulting in 254 evaluated pools. In the mix setting, candidates from all predictors were pooled for each target, resulting in 39 targets with 1,200 candidates per target.

#### Quality-assessment comparison on FoldBench

Because the dedicated quality-assessment methods did not outperform predictor-native confidence measures in the CASP16 analysis, the FoldBench selection analysis was restricted to ranking score, pTM, and pLDDT. Candidate pools contained 200 structures per target from each of the same six predictors. Ground-truth accuracy was measured using TM-score for protein monomers, DockQ for multimeric interfaces, and lDDT-PLI for protein-ligand complexes, following FoldBench original metrics. Candidates lacking either a ground-truth accuracy metric or the corresponding confidence value were excluded. Selection was evaluated in the same single and mix settings. Because predictor performance differs among FoldBench categories, the numbers of evaluated pools and targets are reported separately for each category. Mean Spearman (ρ) and Top-1 ranking loss are reported in Supplementary Fig. 5.

### Computing hardware

Inference was performed on NVIDIA A100 80 GB PCIe and RTX A6000 48 GB GPUs. CPU-only workloads such as MSA and template search were run on dual-socket Intel nodes with 40–48 cores and 188–503 GB of RAM.

## Data Availability

The local MSA and template databases are publicly available from the Hugging Face Hub at https://huggingface.co/datasets/cssbsnu/Thal-Kak_local_db.

Boltz2, Chai-1, ESMFold2, and Protenix v1/v2 predictions and confidence scores for FoldBench are publicly available from Zenodo at https://zenodo.org/records/21947699.

## Code Availability

Thal-Kak is free and open-source software (Apache 2.0), available at https://github.com/CSSB-SNU/Thal-Kak. Google Colab Notebook is available at https://colab.research.google.com/github/CSSB-SNU/Thal-Kak/blob/main/Thalkak.ipynb

## Acknowledgements

This work was supported by the Institute of Information & communications Technology Planning & Evaluation (IITP) [RS-2023-00220628, RS-2025-25442149] and Korea Basic Science Institute(National research Facilities and Equipment Center) [RS-2024-00401698] funded by the Korea government (MSIT).

