## Supplementary Methods, Tables, Figures for "Thal-Kak: unifying biomolecular structure predictors reveals a sampling–selection gap"

##### *S1. Pipeline details*

**Input specification.** A Thal-Kak job is defined by a single input file containing two sections. The *Method* section specifies the alignment method (*msa*), structure predictor or predictors (*structure*), relaxation protocol (*relax*), and seed range (*n_seed*, *seed_start*). The *Entity* section describes the molecular components of the target. Protein, DNA, and RNA entities are specified by sequence and copy number, whereas ligands are specified using SMILES strings[1] or Chemical Component Dictionary (CCD) identifiers[2]. Copy numbers define the stoichiometry of the assembly. The *structure* field can contain multiple predictors, whereas *msa* and *relax* each accept a single method. Supplying multiple methods to either field raises an error rather than silently ignoring additional entries.

Each stage records its effective parameters in a log that is propagated through subsequent stages. Every final model is therefore accompanied by the settings used to generate it. A failure of one predictor is recorded without terminating the remaining prediction jobs, and a pass or fail summary for all requested predictors is reported at the end of the run. The MSA and template search, structure prediction, and relaxation stages can also be executed independently, allowing intermediate files to be reused or replaced with user-provided inputs.

**Structure selection.** Within each predictor's output, candidate structures are ranked by that predictor's native ranking score and screened for polymer bond connectivity. Consecutive residues in polypeptide chains are required to have a C to N distance of no more than 1.5 Å. Consecutive residues in nucleic-acid chains are required to have an O3′ to P distance of no more than 1.8 Å. The five highest-ranking structures that pass these checks are retained as the final prediction set and passed to the configured relaxation stage.

**Notebook interface.** Thal-Kak is additionally distributed as a Jupyter notebook that runs on Google Colab. User-provided sequence, stoichiometry, predictor, relaxation, and seed settings are converted into the same input format used by the command-line workflow, and both interfaces invoke the same underlying pipeline. The notebook provides interactive visualization of selected structures together with available confidence and alignment information. Protein targets are supported directly in the notebook, whereas workflows requiring local nucleic-acid databases are run through the command-line interface.

#### *S2. Local MSA and template search*

##### *S2.1 Search databases*

###### Protein MSA databases

**Overview.** The local protein MSA pipeline uses UniRef[3,4] together with four environmental sequence resources: MGnify[5], Logan[6], BFD[7], and EnVhog[8]. The MMseqs2[9] and HHblits[10] protocols use overlapping but non-identical search collections and database representations. For MMseqs2, resources are represented as expandable sequence-profile databases constructed either from locally generated clusters or from clustering supplied by the source resource. For HHblits, the corresponding resources are represented as HH-suite profile databases. The exact sequence content and clustering therefore differ between the two search protocols, as detailed below and in Supplementary Fig. 6.

**UniRef databases.** For UniRef30, we used the `uniref30_2302` clustered database distributed with ColabFold[11], together with taxonomy and cluster-mapping metadata refreshed in August 2025. UniRef30 was used as distributed and was not reclustered. In the MMseqs2 protocol, the profile generated from UniRef30 is used to seed searches against the environmental collections. UniRef30 is also used for interchain pairing in all local protocols because it provides the NCBI taxonomy annotations[12] required by `mmseqs pairaln`.

The second UniRef source is UniRef100 release `2026_01`, which was clustered locally and prepared for both MMseqs2 and HHblits. UniRef30 and UniRef100 are searched independently rather than treating the newer release as a replacement for UniRef30. Among the sequences accessible through either resource,matched by exact sequence identity, 49.9% are represented by both, 13.8% are accessible through UniRef30 alone, and 36.3% through UniRef100 alone. Searching both therefore provides access to 1.57 times as many sequences as UniRef30 alone (Supplementary Fig. 7).The two resources are combined only when their resulting alignments are merged.

**Environmental sequence resources.** For Logan, we used representative protein sequences released for contigs version 1.0, including SRA accessions available through December 2023. The selected sequences correspond to genes predicted as complete, defined as full-length open reading frames bounded by both a start and a stop codon. Logan distributes human-associated and non-human-associated assemblies separately. Because the non-human collection is substantially larger, it was not constructed as an MMseqs2 profile database and is searched only by the HHblits protocol.

For MGnify, we used the protein cluster representatives from release `2024_04`, which were clustered by the source resource at 90% sequence identity and 90% coverage.

For BFD, the MMseqs2 and HHblits protocols use different representations. The MMseqs2 workflow uses the reduced set of representative sequences, whereas the HHblits workflow uses the complete author-distributed HH-suite database `bfd_metaclust_clu_complete_id30_c90`.

For EnVhog, the MMseqs2 workflow uses the released protein sequences together with the resource's `standard` clustering. The HHblits workflow instead uses the per-family alignments and profile HMMs released at the `enVhog` level. These families are defined by profile-based grouping rather than by the sequence-identity clustering used for the MMseqs2 representation.

**MMseqs2 profile databases.** For locally constructed MMseqs2 resources, source sequences were first converted to MMseqs2 sequence databases using `mmseqs createdb`. Unless stated otherwise, redundant sequences were clustered using `mmseqs linclust` at a minimum pairwise sequence identity of 30% and target coverage of 80% with `--min-seq-id 0.3 -c 0.8 --cov-mode 1`.

Each cluster member was aligned to its representative while retaining the complete alignment backtrace and without applying an E-value cutoff using `mmseqs align -a -e inf`. The resulting alignments were converted to per-cluster sequence profiles and consensus sequences using `mmseqs result2profile` and `mmseqs profile2consensus`. Sequence, header, consensus, and alignment records were packaged into expandable profile databases using `mmseqs tsv2exprofiledb`, and search indices were generated using `mmseqs createindex`.

UniRef100, Logan human, and MGnify were processed using this workflow. BFD used the same profile-construction procedure but was clustered using the cascaded `mmseqs cluster` module at 30% sequence identity and full-length target coverage with `--min-seq-id 0.3 -c 1.0`.

EnVhog was not reclustered. Its published `standard` clustering table was converted into an MMseqs2 cluster database using `mmseqs tsv2db` and subsequently processed through the same profile-construction workflow. UniRef30 was used directly in the representation distributed by ColabFold.

**HH-suite databases.** For HHblits, each database is stored in HH-suite format as three FFindex components containing the per-cluster A3M alignment, profile HMM, and CS219 context-state representation. For the environmental collections, these are independent database builds rather than re-encodings of the MMseqs2 clustering.

MGnify and Logan human and non-human were converted from their clustered member alignments. For each representative cluster, the member alignment was written in A3M format and used to construct the corresponding profile HMM and CS219 representation.

BFD and EnVhog were used from their released profile resources. BFD uses the complete author-distributed HH-suite database `bfd_metaclust_clu_complete_id30_c90`, in contrast to the reduced BFD representation used by MMseqs2. EnVhog uses the per-family A3M alignments and HMMs from the `enVhog` level.

UniRef30 and UniRef100 were converted to HH-suite format using the same source releases described above. The MMseqs2 representation of UniRef30 remains necessary in the HHblits workflow because its taxonomy metadata are used for interchain pairing.

To restrict HMM-HMM comparison to statistically supported profiles, the HMM is not built from clusters whose alignment depth is 50 sequences or fewer, in every database except UniRef100. Clusters whose alignment depth is below three sequences are excluded entirely from every database except UniRef100, so that no component is built for alignments too shallow to yield a meaningful profile.

**Template database**

Templates are searched against a snapshot of the Protein Data Bank[13] containing entries released through 24 February 2026. Protein sequence records were deduplicated by PDB identifier, author chain identifier, and sequence. This removes repeated chain representations arising from different assemblies, structural models, or alternate-location variants. The resulting search collection contains 1,025,280 chains from 244,541 PDB entries and 178,610 distinct protein sequences. No additional sequence clustering or redundancy filtering is applied. Coordinates are stored as one compressed mmCIF file per PDB entry and extracted when a template hit is retained.

**RNA MSA databases**

RNA MSAs are generated using two nucleotide sequence resources: the Rfam family sequence set (`Rfam.fa`)[14] and the active RNAcentral sequence set[15], both downloaded in January 2026.

Each resource was clustered at 90% sequence identity and 80% target coverage using `--min-seq-id 0.9 -c 0.8 --cov-mode 1`. Rfam was clustered using `mmseqs easy-cluster`. Because of its larger size, RNAcentral was clustered using the linear-time `mmseqs easy-linclust` implementation.

The representative sequences from each resource were converted to binary `nhmmer`[16] sequence databases using `makehmmerdb` and indexed for sequence retrieval using `esl-sfetch --index`.

##### *S2.2 MSA search protocols*

**Overview.** Three local protein search protocols are provided: MMseqs2, HHblits, and a combined protocol that executes both. All three produce the same downstream alignment representation, consisting of one ColabFold complex-format A3M per target together with per-chain paired and unpaired alignments. Unless stated otherwise, the search topology and parameter settings follow the ColabFold workflow[11].

**MMseqs2 protocol.** The MMseqs2 search topology follows the ColabFold MSA server. Each distinct protein sequence is first searched against UniRef30 using `mmseqs search --num-iterations 3 -s 8.0 -e 0.1 --max-seqs 10000 --k-score seq:96,prof:80 -a`.

Accepted hits are expanded to their cluster members using `mmseqs expandaln --expansion-mode 0 -e inf --expand-filter-clusters 1 --max-seq-id 0.95`. Expanded hits are realigned against the iteration-3 profile using `mmseqs align -e 10 --max-accept 100000 --alt-ali 10 -a`, filtered using `mmseqs filterresult --qid 0 --qsc 0.8 --diff 0 --max-seq-id 1.0 --filter-min-enable 100`, and written to the final alignment using `mmseqs result2msa --msa-format-mode 6 --filter-msa 1 --diff 3000 --qid 0.0,0.2,0.4,0.6,0.8,1.0 --qsc 0 --max-seq-id 0.95 --filter-min-enable 1000`.

UniRef100 is searched independently using the same search procedure.

The profile derived from the UniRef30 search is subsequently used to search each environmental database independently. Environmental hits are expanded using `mmseqs expandaln --expansion-mode 0 -e inf`, which admits all cluster members associated with an accepted hit. Searching the environmental collections separately preserves database-level provenance and allows individual resources to be added or removed without constructing a pre-merged database.

**HHblits protocol.** Each distinct protein sequence is first searched against UniRef100 using `hhblits -n 3 -e 0.001 -realign_max 100000 -maxfilt 100000 -min_prefilter_hits 1000 -p 20 -Z 500`.

The resulting A3M alignment is then used as the query for a separate search of each remaining databases (UniRef30 2302, complete BFD, MGnify, Logan human, Logan non-human, and EnVhog) using the same settings. Every search starts from this same alignment, and the profile is refined within the three iterations of each search rather than across databases. The relaxed prefilter, maximum-filter, and realignment settings follow the HHblits invocation used in AlphaFold3.

The HH-suite representation of UniRef100 does not retain the NCBI taxonomy identifiers required for interchain pairing. For targets containing multiple distinct protein sequences, the HHblits workflow therefore generates the paired alignment separately using the MMseqs2 pairing procedure against UniRef30 described below.

**Combined protocol.** The MMseqs2 and HHblits workflows are executed independently to completion. Their per-chain alignments are then merged by database class rather than by search engine, so that redundancy filtering is applied once to each class.

Within a database class, MMseqs2-derived rows are placed before HHblits-derived rows. Exact duplicate sequences are removed while retaining the first occurrence. Consequently, when both engines recover the same sequence, the MMseqs2 representation is retained. This ordering also gives MMseqs2-derived rows priority if the merged alignment exceeds the final depth limit.

**Per-chain merging.** Per-database alignments for each chain are grouped into UniRef-derived and environmental sequence sets and concatenated within each group in a fixed database order. Sequences with identical residue content after removal of gaps and conversion of insertion residues to uppercase are collapsed while retaining the first occurrence.

Each group is initially filtered using hhfilter (`hhfilter -id 90 -cov 75`). If fewer than 2,000 sequences remain, the group is instead filtered with lower coverage threshold (`hhfilter -id 90 -cov 50`). If fewer than 100 sequences remain after this second filtering step, the unfiltered but deduplicated group is retained.

The resulting UniRef-derived and environmental alignments are concatenated, deduplicated again across groups, and truncated to the final depth budget. The resulting alignment depth is therefore determined by sequence redundancy and coverage rather than by a fixed quota assigned to each source database. The same merging procedure is used for all three local search protocols.

**Interchain pairing.** For targets containing more than one distinct protein sequence, an interchain-paired alignment is constructed so that corresponding rows across chains originate from the same species. Pairing is performed against UniRef30 in all local search protocols because it provides the NCBI taxonomy annotations required by `mmseqs pairaln`. Neither UniRef100 nor the environmental databases contribute directly to the paired block.

The paired alignment is generated using the same initial `mmseqs search` invocation as in the MMseqs2 protocol. Accepted hits are expanded using `mmseqs expandaln --expansion-mode 0 -e inf --expand-filter-clusters 0 --max-seq-id 0.95` and realigned against the profile using `mmseqs align -e 0.001 --max-accept 1000000`.

An initial pairing step is performed using `mmseqs pairaln --pairing-mode 0 --pairing-dummy-mode 0`. The resulting records are back-aligned using `mmseqs align -e inf -a`, followed by a second pairing step with `mmseqs pairaln --pairing-mode 0 --pairing-dummy-mode 1`. The second step introduces gap-only rows when a species contains a homolog for one chain but not another, ensuring that corresponding paired blocks contain the same number of rows for all chains. The final paired alignment is written using `mmseqs result2msa --msa-format-mode 6`.

**Assembly and depth budget.** Paired and unpaired per-chain alignments are assembled into one ColabFold complex-format A3M. The header records the sequence length and copy number of each entity. The paired block concatenates corresponding rows across chains, whereas the unpaired block contains single-chain sequences padded with gaps to the full complex length.

For each chain, the paired block is capped at 8,192 records. The total alignment depth is capped at 16,384 records including the query, with the remaining capacity assigned to unpaired sequences.

**RNA MSA search.** Each RNA query is searched against both RNA sequence databases described in Section S2.1 using `nhmmer -E 0.001 --incE 0.001 --rna --watson --F3 0.00005`, following the AlphaFold3 parameterization. Queries shorter than 50 nucleotides use `--F3 0.02`.

Up to 10,000 hits are retained from each database. Hits are pooled across resources and deduplicated by exact sequence. The pooled sequences are then realigned to a profile HMM constructed from the query using `hmmbuild --rna` and `hmmalign --rna --mapali`. The resulting Stockholm alignment is converted to A3M format.

##### *S2.3 Template search protocols*

Template search is paired with the protein MSA protocol used for the target. The combined MSA protocol executes both template-search procedures and merges the resulting hits.

**MMseqs2 template search.** The UniRef30-derived profile generated during the MMseqs2 MSA search is reused as the query and searched against the local template sequence database using `mmseqs search -s 7.5 -e 0.1 -a`.

Alignments are converted to tabular form using `mmseqs convertalis --format-output query,target,fident,alnlen,mismatch,gapopen,qstart,qend,tstart,tend,evalue,bits,cigar`.

**HMMER template search.** For the HHblits workflow, the UniRef100 alignment generated in the first search stage is reduced to its query-match columns and converted into a profile HMM[17] using `hmmbuild --hand --amino`. A reference annotation (`#=GC RF`) is used so that profile match states correspond to query positions.

The resulting profile is searched against the template FASTA using `hmmsearch --noali --F1 0.1 --F2 0.1 --F3 0.1 -E 100 --incE 100 --domE 100 --incdomE 100`, following the AlphaFold3 parameterization[18].

Template hits are required to contain at least 10 aligned residues, cover at least 10% of the query, and have a fraction of query-identical residues no greater than 0.95, following the AlphaFold3 template-filtering criteria.

**Date restriction.** The local template search supports a user-specified release-date cutoff for applications that require temporal control of template availability. Template availability is defined using the earliest `_pdbx_audit_revision_history.revision_date` recorded in the corresponding mmCIF file, with the deposition date used when revision-history information is unavailable. Entries released after the specified cutoff are excluded. Using the release date rather than the deposition date also excludes structures that were deposited before the cutoff but not publicly released until afterward.

**Output.** For both template search engines, hits are ranked by E-value, deduplicated, and capped at 20 templates per chain, of which the four best per chain are passed to the predictors. Each retained hit is represented by one row in a 13-column tabular hit file containing the query-to-template residue correspondence encoded as a CIGAR string, together with the corresponding mmCIF structure extracted from the local PDB snapshot.

##### *S2.4 Search software and configuration provenance*

Local database construction and sequence searches use MMseqs2 release 18-8cc5c, HH-suite version 3.3.0, and HMMER version 3.4.0.

MMseqs2 is used for construction of MMseqs2 protein databases, protein MSA searches, interchain pairing, and MMseqs2-based template searches. HH-suite is used for HHblits database construction, HHblits searches, and per-chain alignment filtering. HMMER and Easel are used for RNA database construction, RNA sequence searches, RNA profile construction and alignment, and HMM-based template searches.

All search parameters and database selections are read from protocol-specific configuration files. The effective configuration used for each alignment is recorded with the corresponding Thal-Kak output.

#### *S3. Structure prediction details*

**Shared input and predictor-specific backends.** Each predictor-specific runner combines the shared Thal-Kak job description with its own configuration and converts these inputs into the native format required by the corresponding backend. Alignments, templates, ligands, and assembly stoichiometry are supplied from the same upstream representation, subject to the input features supported by each predictor. Templates and RNA alignments are therefore passed only to backends that accept them. ESMFold2[19] can be run with or without an external alignment.

Predictor-specific configuration files define inference settings including the random seed range, number of recycles, number of diffusion steps, number of samples per seed, and handling of MSA depth. These configurations also expose predictor-specific MSA subsampling or truncation where applicable. The complete alignment can be supplied up to the featurization limit of a predictor when that behavior is supported by the backend. The effective predictor configuration is recorded with each run.

**Sampling and output.** Predictions are generated over a contiguous range of random seeds using each backend's native stochastic sampling interface. No perturbation is applied to the upstream MSA, template, or molecular inputs to induce additional diversity.

For each generated candidate, the confidence measures reported by the predictor are collected into a table with a common schema. These include mean pLDDT, predicted TM-score, interface predicted TM-score, and the predictor's composite ranking score where available. Per-residue pLDDT and predicted aligned error are retained when provided by the corresponding backend. Standardized structures, confidence summaries, and plots are collected in a common output directory, while each predictor's native output layout is preserved alongside them.

#### *S4. Restrained relaxation*

**Pre-relaxation validation.** Before relaxation, prochiral methyl labels are standardized for Val CG1 and CG2 atoms and Leu CD1 and CD2 atoms without altering their coordinates. Protein C-terminal carboxylate groups are evaluated using C to O bond lengths, bond angles, and planarity. Missing or distorted O or OXT atoms are reconstructed as planar sp2 carboxylates with C to O bond lengths of 1.25 Å and bond angles of approximately 120 degrees.

**Restrained energy minimization.** Validated structures are subjected to restrained all-atom energy minimization using OpenMM 8.4.0[20] and PDBFixer 1.12.0[21] with a modified AlphaFold2 relaxation procedure[7]. Missing atoms and hydrogens are added assuming pH 7.0. Explicit water molecules are removed, and solvent is represented using the OBC2 implicit solvent model[22].

The force-field stack comprises amber19-all with ff19SB for proteins[23], OL21 for DNA[24], and OL3 for RNA[25], together with GLYCAM_06j-1 for carbohydrates[26], monatomic-ion parameters, and GAFF 2.11 with AM1-BCC charges for small-molecule ligands[27,28]. All non-hydrogen atoms are restrained relative to their starting coordinates using confidence-dependent flat-bottom harmonic potentials.

Up to three minimization cycles are performed, with a maximum of 2,000 L-BFGS iterations in each cycle[29]. Residues with newly detected stereochemical violations are released from positional restraints in subsequent cycles. Initial and final potential energies are recorded in kcal/mol. Models are retained only when minimization completes successfully.

#### *S5. Additional benchmark implementation details*

**Handling of modified residues in DockQ.** DockQ v2.1.3[30] can classify polymer chains containing modified residues as small molecules, which results in truncation of the reconstructed polymer sequence. For affected structures, polymer records corresponding to standard residues were restored and each chain sequence was reconstructed residue by residue using the AlphaFold3 CCD one-letter mapping before DockQ evaluation. The same corrected chain representation was used together with the chain correspondence established by OpenStructure[31].

**Predictor-specific inference settings.** Thal-Kak stores inference parameters separately for each predictor so that differences in backend configuration remain explicit. These configurations specify the random seeds, number of candidate structures generated per seed, recycling and diffusion settings where applicable, and predictor-specific handling of MSA depth. The exact configuration files used to generate the benchmark prediction pools are distributed with the analysis code.

**Output provenance.** Candidate structures retain the identity of the predictor, seed, and within-seed sample from which they were generated. Predictor-native confidence measures are preserved without cross-model normalization. This metadata is propagated to the pooled candidate sets used for seed-budget and quality-assessment analyses.

##### *S6. Local MSA search benchmark*

**Target sets**. The alignment comparison in Supplementary Fig. 2b,c uses target sets separate from those of the sampling and selection benchmarks. Monomers were drawn from the 334 protein-monomer targets of FoldBench[32] Supplementary Table 3, retaining those for which at least one of AlphaFold 3[18], Chai-1[33] or Protenix[34] scored below TM-score 0.7 in the FoldBench runs. Multimers were drawn from the 227 bioassemblies of the FoldBench protein-protein interaction set. Both sets were restricted to targets released on or after 1 January 2024, leaving 9 monomers and 95 multimers, the latter after excluding 8 targets whose input composition disagreed with the reference biological assembly.

**Predictions**. Both arms were run template-free through AlphaFold 3 with five random seeds and five candidate structures per seed, giving 25 candidates per target in each arm. The Thal-Kak arm used the combined local protocol, in which the MMseqs2 and HHblits alignments are merged (Supplementary Methods S2.2); the baseline arm used the ColabFold API alignment[11]. Accuracy in Supplementary Fig. 2c is reported for the oracle candidate of each pool.

#### Supplementary Table

**Supplementary Table 1.** Statistical similarity between Thal-Kak benchmark scores and FoldBench reference values. We performed a one-sided Wilcoxon signed-rank non-inferiority test (margin 0.05, $\alpha = 0.05$) to demonstrate that Thal-Kak results are not statistically inferior to those from FoldBench. Analysis compares the highest-ranking-score decoy (among five seeds) for each target against the corresponding FoldBench reference value. *n* denotes the number of targets, pooled across AF3, Boltz-2, and Chai-1.

| **Target type** | *n* | **Wilcoxon** *p* |
|---|---|---|
| Protein–Protein | 366 | $4.6 \times 10^{-43}$ |
| Antibody–Antigen | 208 | $9.9 \times 10^{-21}$ |
| Protein–Peptide | 79 | $5.0 \times 10^{-4}$ |
| Protein–NA | 602 | $5.9 \times 10^{-55}$ |
| Protein–Ligand | 574 | $2.6 \times 10^{-47}$ |
| Monomer (protein) | 466 | $3.6 \times 10^{-74}$ |
| All targets (pooled) | 2295 | $8.1 \times 10^{-226}$ |

### Supplementary Figures

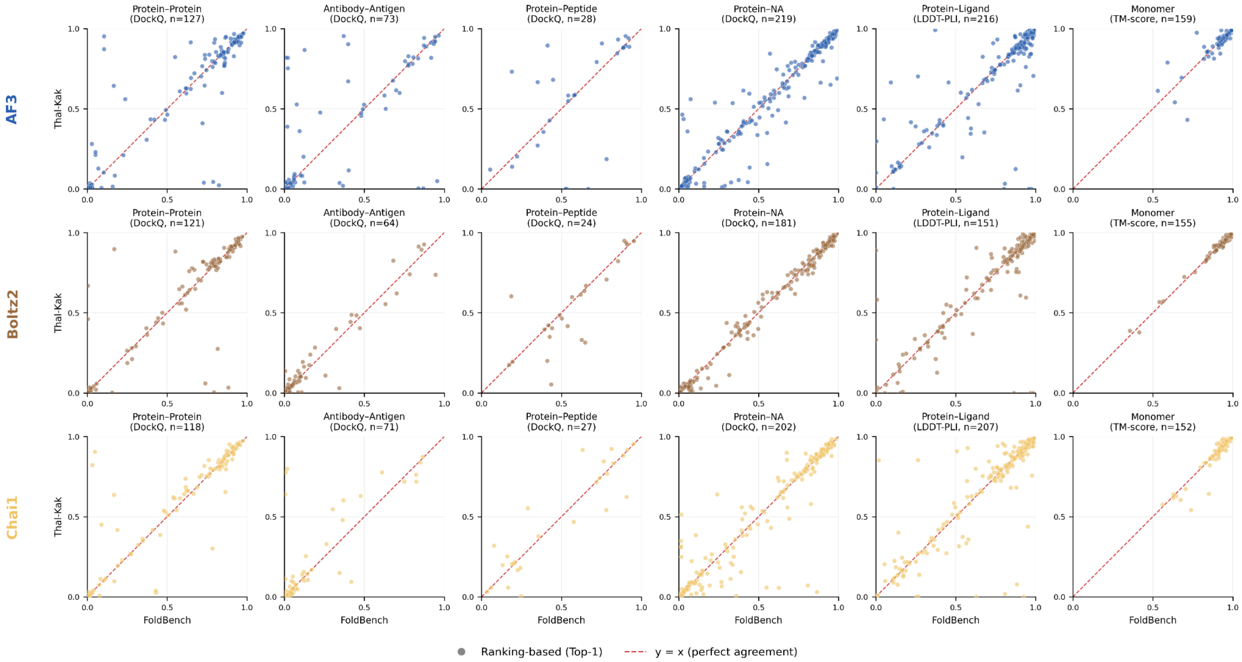


**Supplementary Figure 1. Reproduction of FoldBench evaluation scores.** Per-target agreement between structural accuracy scores reproduced with the Thal-Kak evaluation pipeline (using ColabFold API MSA) and those reported by FoldBench (using each method-specific MSA pipeline). Each point represents one target, evaluated using the candidate with the highest predictor-native ranking score among five randomly sampled seeds. Rows correspond to AF3, Boltz-2, and Chai-1, and columns to the six FoldBench target classes. The evaluation metric and number of targets are indicated above each panel. The red dashed line indicates perfect agreement (y=x).

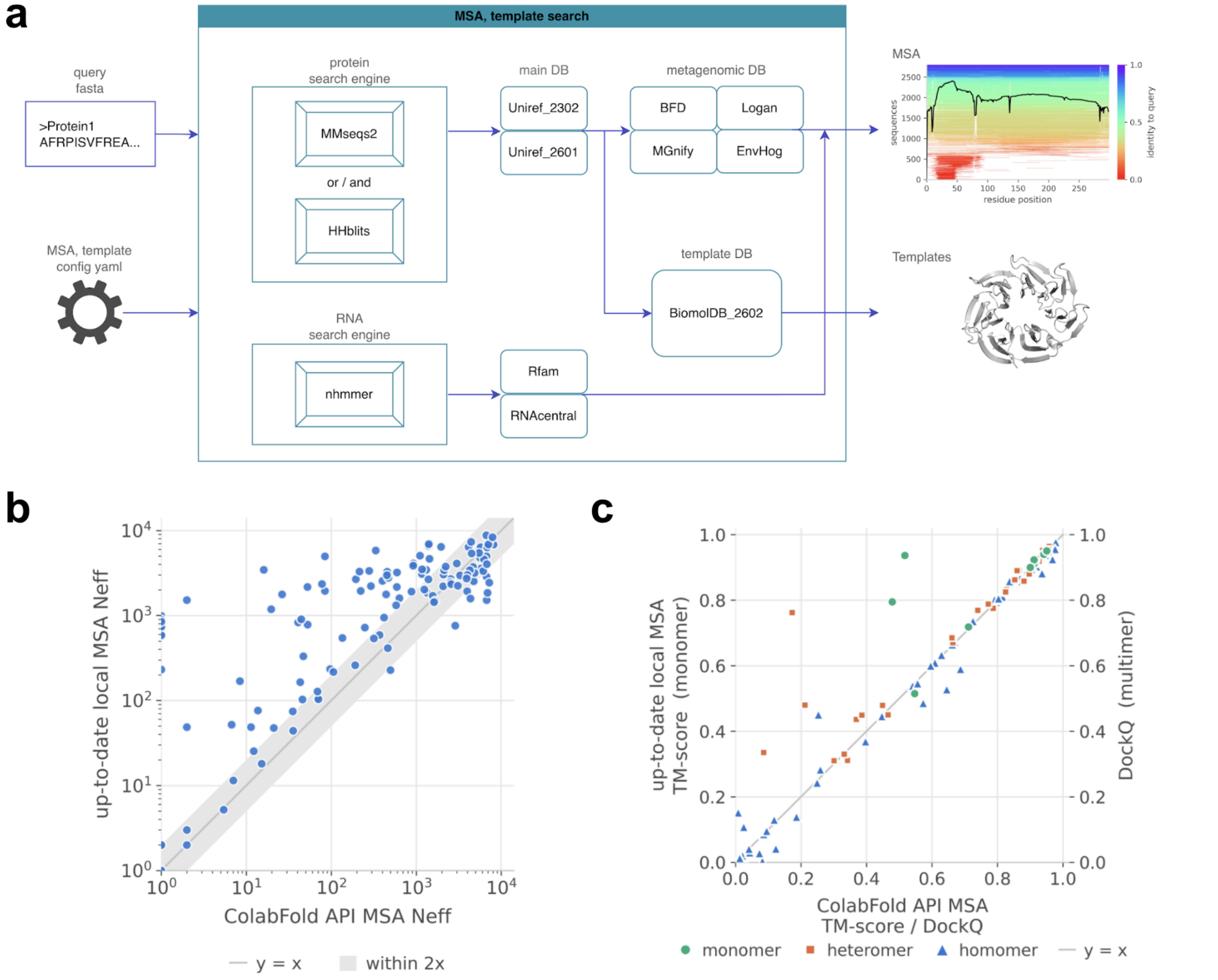


**Supplementary Figure 2. Thal-Kak's local MSA and template search. a,** Overview of the local search workflow. Protein sequences are searched using MMseqs2, HHblits, or both against UniRef30/100 and environmental sequence databases including BFD, Logan, MGnify, and EnVhog. RNA sequences are searched with nhmmer against Rfam and RNAcentral. Templates are retrieved from the PDB snapshot (February 2026). **b,** Effective MSA depth, measured by Neff, obtained with the Thal-Kak local search using MMseqs2 and HHblits versus the ColabFold API. The diagonal indicates equal depth and the shaded region indicates values within twofold. **c,** Structure-prediction accuracy using local versus ColabFold-API alignments. TM-score is shown for monomers and DockQ for homomeric and heteromeric complexes. The diagonal indicates equal performance. For the benchmark details, see Supplementary Methods S6.

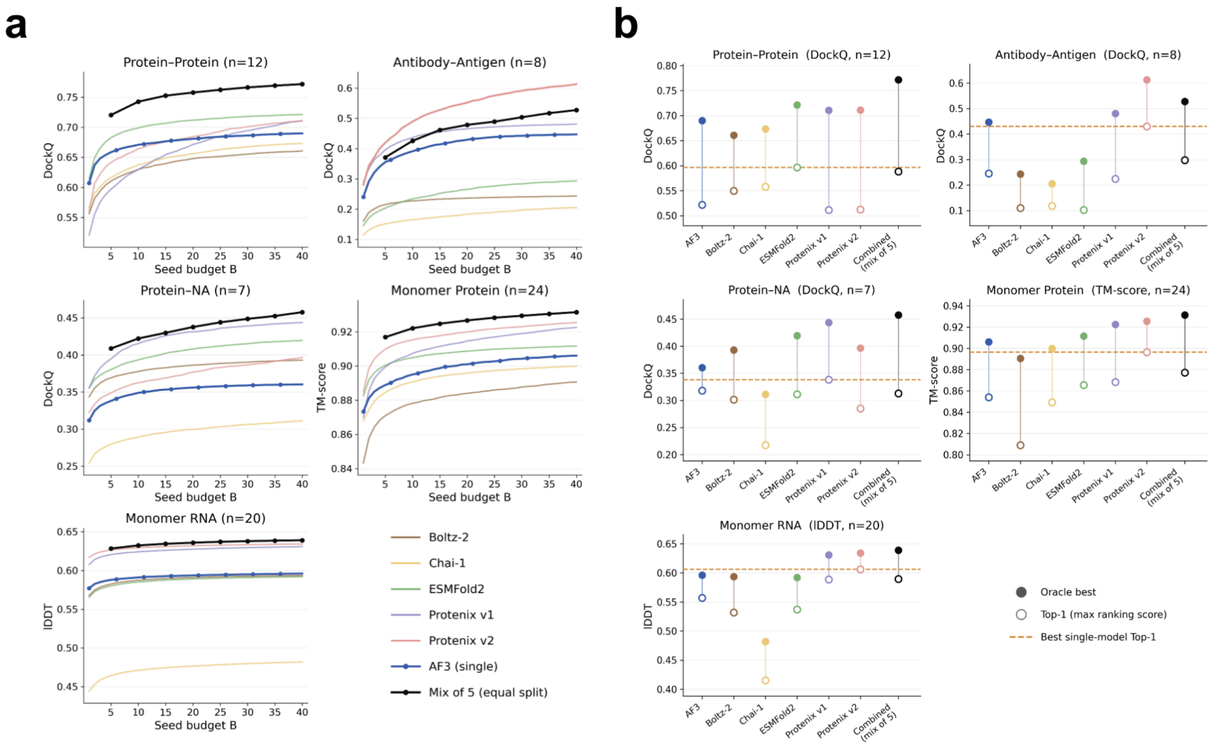


**Supplementary Figure 3. Multi-predictor sampling and selection on CASP16. a,** Oracle sampling performance across five CASP16 target classes as a function of total seed budget B. AF3 is shown in blue, the equal-split mixture of the five released predictors in black, and individual released predictors as lighter curves. The mixture assigns B/5 seeds to each predictor. Performance is measured by DockQ for protein-protein, antibody-antigen, and protein-NA targets, TM-score for protein monomers, and lDDT for RNA monomers. Curves show mean performance across targets and repeated random seed subsampling. **b,** Oracle and Top-1 performance at a fixed budget of 40 seeds. Filled circles indicate the best sampled structure and open circles the structure selected by the predictor-native ranking score. Single predictors use 40 seeds, whereas the five-model mixture uses 8 seeds per predictor. The dashed line marks the best Top-1 performance among individual predictors.

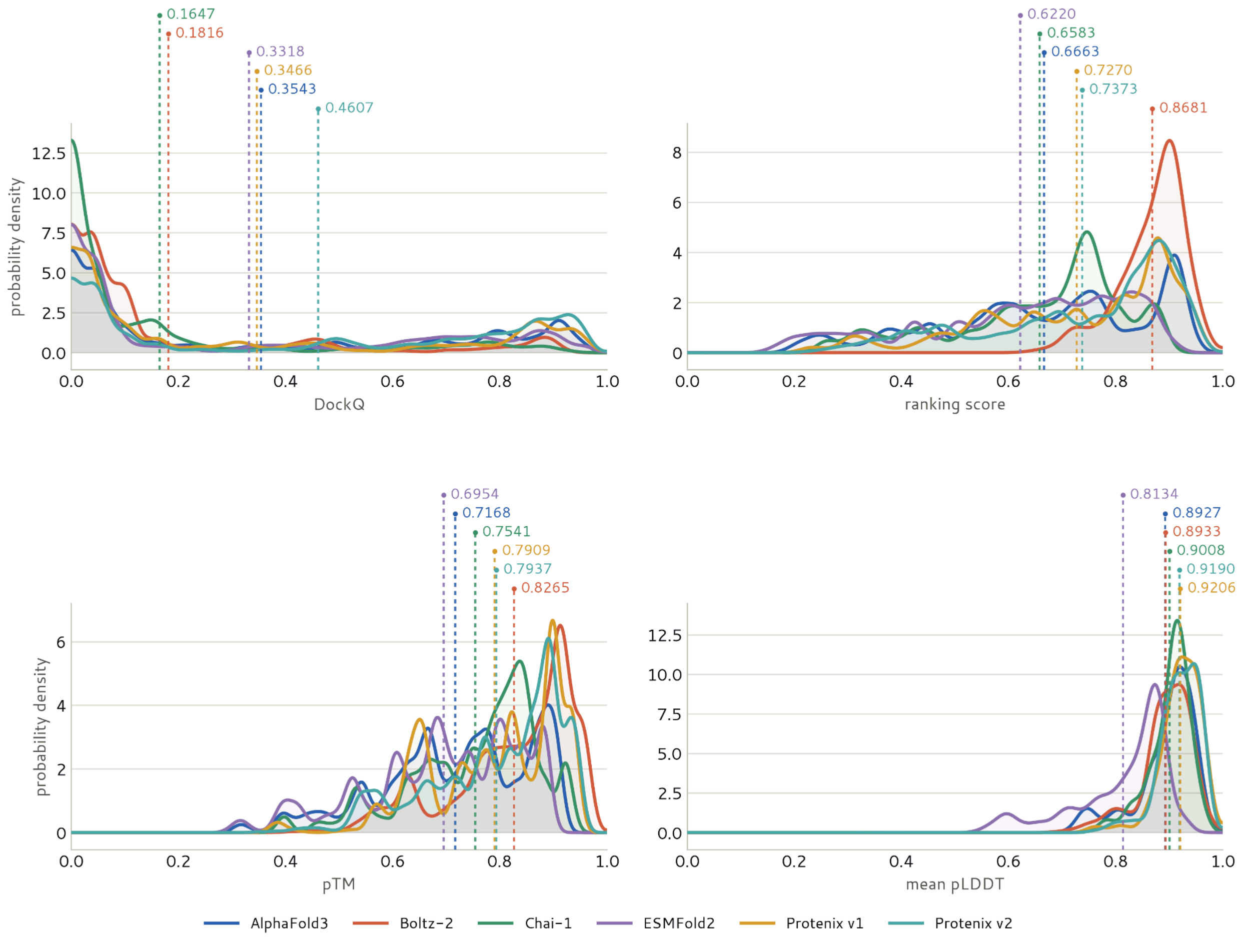


**Supplementary Figure 4. Cross-predictor differences in confidence distributions for FoldBench antibody-antigen targets.** Probability density distributions of ground-truth DockQ, predictor-native ranking score, pTM, and mean pLDDT for structures generated by AlphaFold3, Boltz-2, Chai-1, ESMFold2, Protenix v1, and Protenix v2. Dashed vertical lines indicate the mean value for each predictor, with the corresponding values shown above. Negative ranking scores are excluded from the ranking-score distribution.

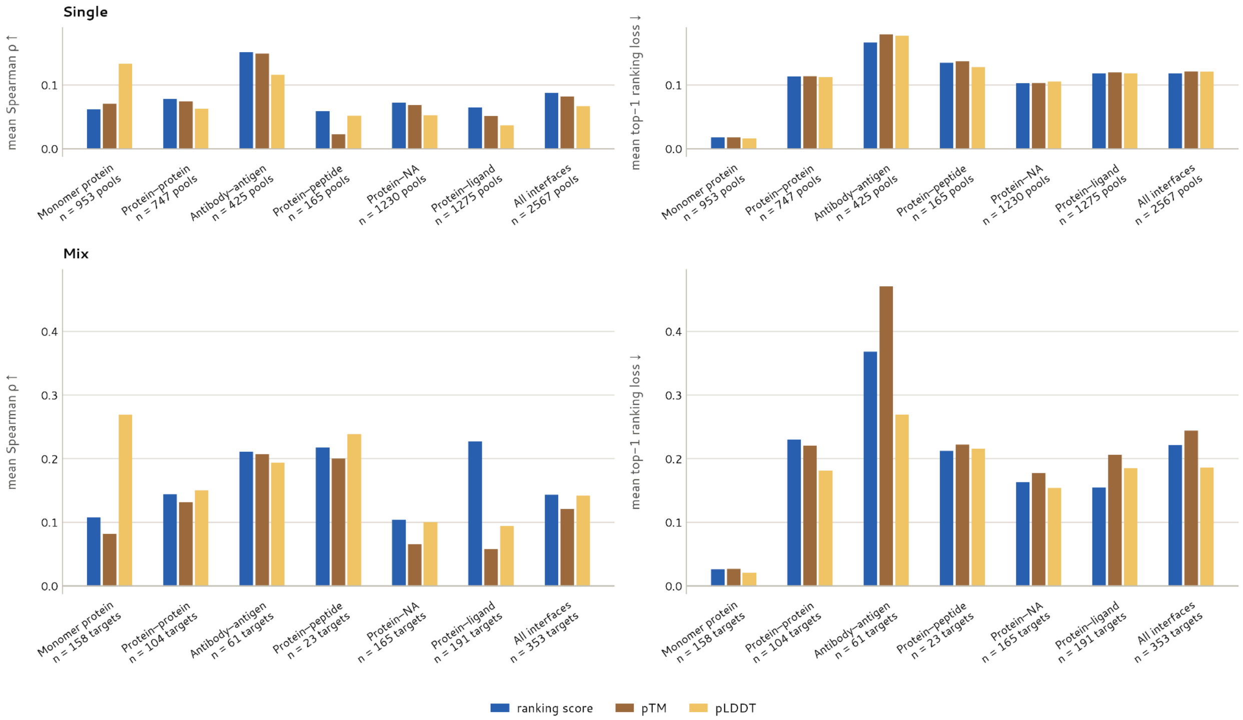


**Supplementary Figure 5. Confidence-based structure selection on FoldBench.** Performance of predictor-native ranking score, pTM, and pLDDT for structure selection across FoldBench target classes. The top panels evaluate candidates from one predictor at a time (Single), whereas the bottom panels pool the candidates of all six predictors for each fully covered target (Mix). Ground-truth accuracy is defined using the corresponding FoldBench metric for each target class. Selection performance is measured by mean Spearman correlation between confidence and accuracy, with higher values indicating better ranking, and mean Top-1 ranking loss, with lower values indicating better selection. Numbers below each category indicate the evaluated pools or targets.

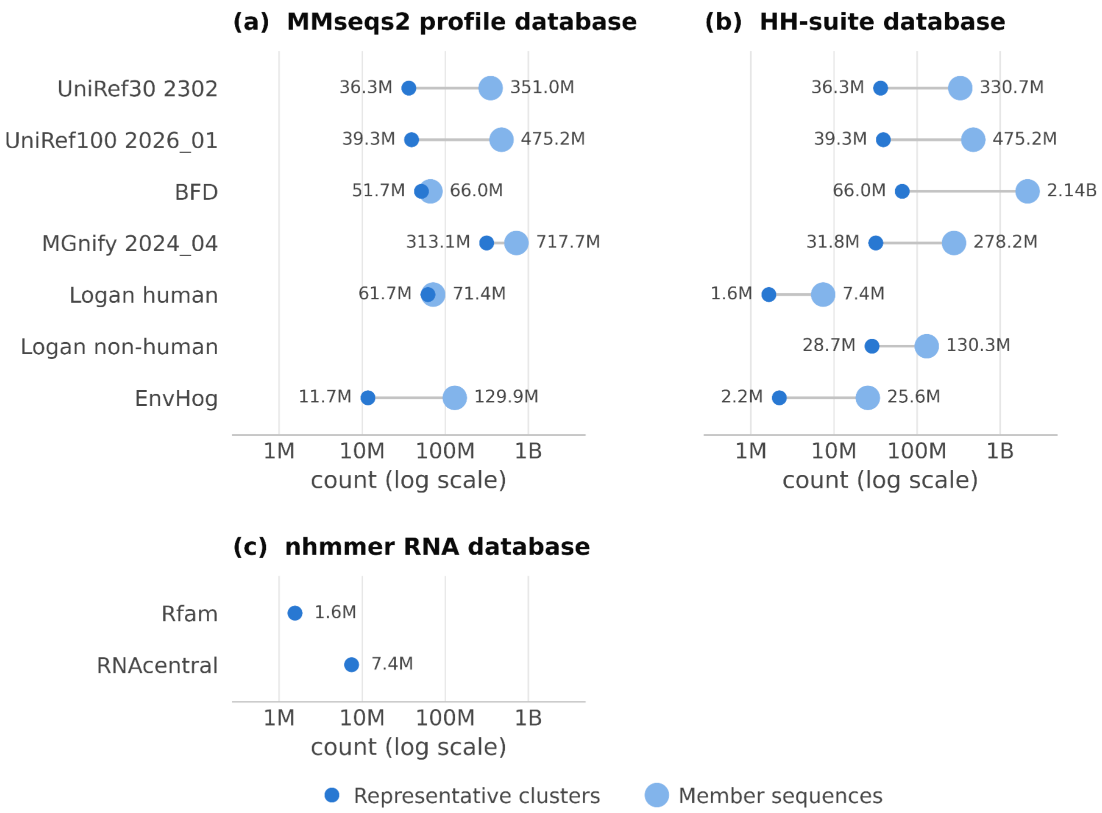


**Supplementary Figure 6**. **Sequence collections used by the local MSA search protocols.** **a,** MMseqs2 profile databases. **b,** HH-suite databases used by the HHblits protocol. **c,** RNA sequence databases used by the nhmmer protocol. For each collection, dark circles indicate the number of representative clusters and light circles the number of member sequences in the underlying collection, plotted on a logarithmic scale. In the protein databases, member sequences can contribute to the final alignment through cluster expansion in MMseqs2 or stored cluster alignments in HH-suite. For the RNA databases, nhmmer searches only the representative sequences.

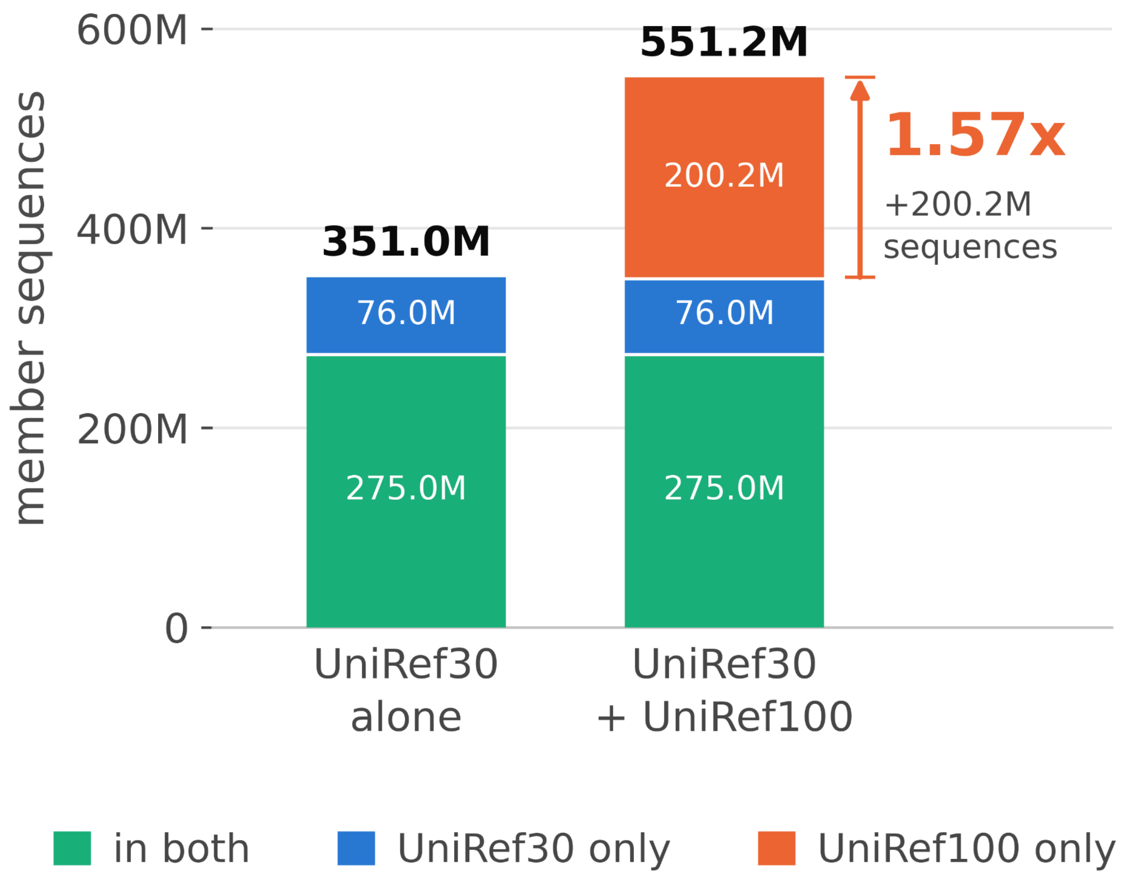


**Supplementary Figure 7. Sequence coverage gained by adding UniRef100 to UniRef30.** Member sequences accessible when searching UniRef30 2302 alone or together with UniRef100 2026_01. Green indicates sequences shared by both databases, blue sequences unique to UniRef30, and orange sequences unique to UniRef100. Adding UniRef100 increases the accessible sequence pool from 351.0 million to 551.2 million sequences, a 1.57-fold increase. Of the combined pool, 275.0 million sequences are shared between the two databases. Sequences were matched by exact identity of their residue strings in the MMseqs2 builds: all 826 million member sequences of the two databases were reduced to 64-bit digests and the two digest sets intersected.
